# UnCOT-AD: Unpaired Cross-Omics Translation Enables Multi-Omics Integration for Alzheimer’s Disease Prediction

**DOI:** 10.1101/2024.10.28.620592

**Authors:** Abrar Rahman Abir, Sajib Acharjee Dip, Liqing Zhang

**Affiliations:** Department of Computer Science and Engineering, Bangladesh University of Engineering and Technology, Dhaka-1000, Bangladesh; Department of Computer Science, Virginia Polytechnic Institute and State University, Blacksburg, 24060, VA, United States

**Keywords:** Alzheimer’s Disease, Multi-omics Integration, Cross Omics Translation

## Abstract

Alzheimer’s Disease (AD) is a progressive neurodegenerative disorder, posing a growing public health challenge. Traditional machine learning models for AD prediction have relied on single omics data or phenotypic assessments, limiting their ability to capture the disease’s molecular complexity and resulting in poor performance. Recent advances in high-throughput multi-omics have provided deeper biological insights. However, due to the scarcity of paired omics datasets, existing multi-omics AD prediction models rely on unpaired omics data, where different omics profiles are combined without being derived from the same biological sample, leading to biologically less meaningful pairings and causing less accurate predictions. To address these issues, we propose **UnCOT-AD**, a novel deep learning framework for **Un**paired **C**ross-**O**mics **T**ranslation enabling effective multi-omics integration for **AD** prediction. Our method introduces the first-ever cross-omics translation model trained on unpaired omics datasets, using two coupled Variational Autoencoders and a novel cycle consistency mechanism to ensure accurate bidirectional translation between omics types. We integrate adversarial training to ensure that the generated omics profiles are biologically realistic. Moreover, we employ contrastive learning to capture the disease specific patterns in latent space to make the cross-omics translation more accurate and biologically relevant. We rigorously validate UnCOT-AD on both cross-omics translation and AD prediction tasks. Results show that **UnCOT-AD** empowers multi-omics based AD prediction by combining real omics profiles with corresponding omics profiles generated by our cross-omics translation module and achieves state-of-the-art performance in accuracy and robustness. Source code is available at https://github.com/abrarrahmanabir/UnCOT-AD

## 1 Introduction

Alzheimer’s Disease (AD) is a progressive neurodegenerative disorder that primarily affects the elderly and is characterized by cognitive decline, memory loss, and ultimately, a loss of bodily functions. The number of AD cases is expected to increase significantly in the coming decades, creating a serious public health challenge globally [1]. Despite numerous studies attempting to identify molecular risk factors involved in AD pathogenesis, the precise mechanisms underlying AD occurrence and progression remain poorly understood. Current treatments for AD can only alleviate symptoms without addressing the root causes of the disease [2,3].

Research has primarily focused on analyzing phenotypic data, such as magnetic resonance imaging (MRI) and neuropsychological assessments [4]. Biomarkers associated with AD pathology, such as *β*-amyloid deposition and tau proteins, have been explored as well, with some recent studies incorporating these biomarkers for more accurate diagnoses [5,6]. However, a significant limitation in many existing AD prediction models is their reliance on single omics, rather than integrating multi-omics data to capture the complexity of AD. Recent advances have allowed for the collection of multi-omics data, which provide a more comprehensive view of biological systems [7,8,9,10,11]. A few studies proposed methods that utilize multi-omics data for prediction. For example, gene expression and DNA methylation data were combined to predict AD [12,13,6]. As paired multi-omics data from the same group of people is not available, the studies used all possible pairs of gene expression profiles from one group of people and DNA methylation profiles from another group of people as surrogate of paired data to predict AD. Using all possible pairs of gene expression and DNA methylation profiles can introduce biologically irrelevant and unrealistic combinations, which adversely affects prediction performance.

To address the challenges posed by the scarcity of paired multi-omics data and the limitations of current multi-omics based AD prediction methods, we propose a novel deep learning framework, **UnCOT-AD** for **Un**paired **C**ross-**O**mics **T**ranslation and multi-omics integration for **AD** prediction. UnCOT-AD performs cross-omics translation using unpaired training datasets, unlike state-of-the-art models such as BABEL [14] and Polarbear [15] that require paired data. Our major contributions are:

1. **Cross-Omics Translation From Unpaired Data:** We introduce a novel Cross-Omics Translation module to perform cross-omics translation using unpaired omics datasets. To the best of our knowledge, this is the first work of cross-omics translation trained on unpaired data. This method allows us to map between different omics types, such as gene expression and DNA methylation, and generate one omics profile from another, even when direct pairings between the two omics types are not available while training. This approach addresses a significant gap in multi-omics integration by enabling biologically meaningful data generation across modalities.
2. **Multi-Omics Based AD Prediction Using Generated Paired Omics Data :** Using the cross-omics translation module, we are able to perform multi-omics-based AD prediction even in the absence of fully paired datasets. By generating a corresponding omics profile (e.g., DNA methylation) from real omics data (e.g., gene expression), we effectively create paired multi-omics data and then fuse the two modalities with our prediction module for AD prediction.

Additionally, as our cross-omics translation module is designed to be compatible with any two omics types and our translation and prediction modules are separate, our method can be applied solely for cross-omics translation between two modalities, even in the absence of paired data. We rigorously validate UnCOT-AD on both cross-omics translation performance and AD prediction tasks, achieving state-of-the-art results in accuracy and robustness.

## 2 Methodology

Our method is divided into two steps. In the first step, a Cross-Omics Translation Module is trained on unpaired omics data, meaning there is no one-to-one correspondence between samples from two different types of omics datasets. The translation module learns a bidirectional mapping such that, at inference, it can take one omics profile as input and translate it to the corresponding profile in the other omics. Next, a Prediction Module combines the real omics profile and the translated omics profile, predicted from the real omics using the pretrained translation module, to predict AD.

### 2.1 Cross-Omics Translation Module

The goal of Cross-Omics Translation Module is to learn bidirectional mappings between two omics data types, denoted as 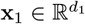 and 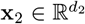 where the given training samples are unpaired (Figure 1). *d*_1_ and *d*_2_ represent the respective dimensionalities of the two omics data types. The empirical data distributions of the two types are denoted as **x**_1_ ∼ *P*_data1_(**x**_1_) and **x**_2_ ∼ *P*_data2_(**x**_2_). We employ two separate Variational Autoencoders (VAEs)[16]. Despite being unpaired, to ensure consistent translations between the two modalities, we incorporate a cycle consistency mechanism. Additionally, three adversarial discriminators are utilized to enforce modality invariance in the latent space and generate biologically realistic omics profiles. To capture AD-specific features, a contrastive loss is introduced to encourage the separation of AD and control samples in the latent space.

**Fig. 1:**
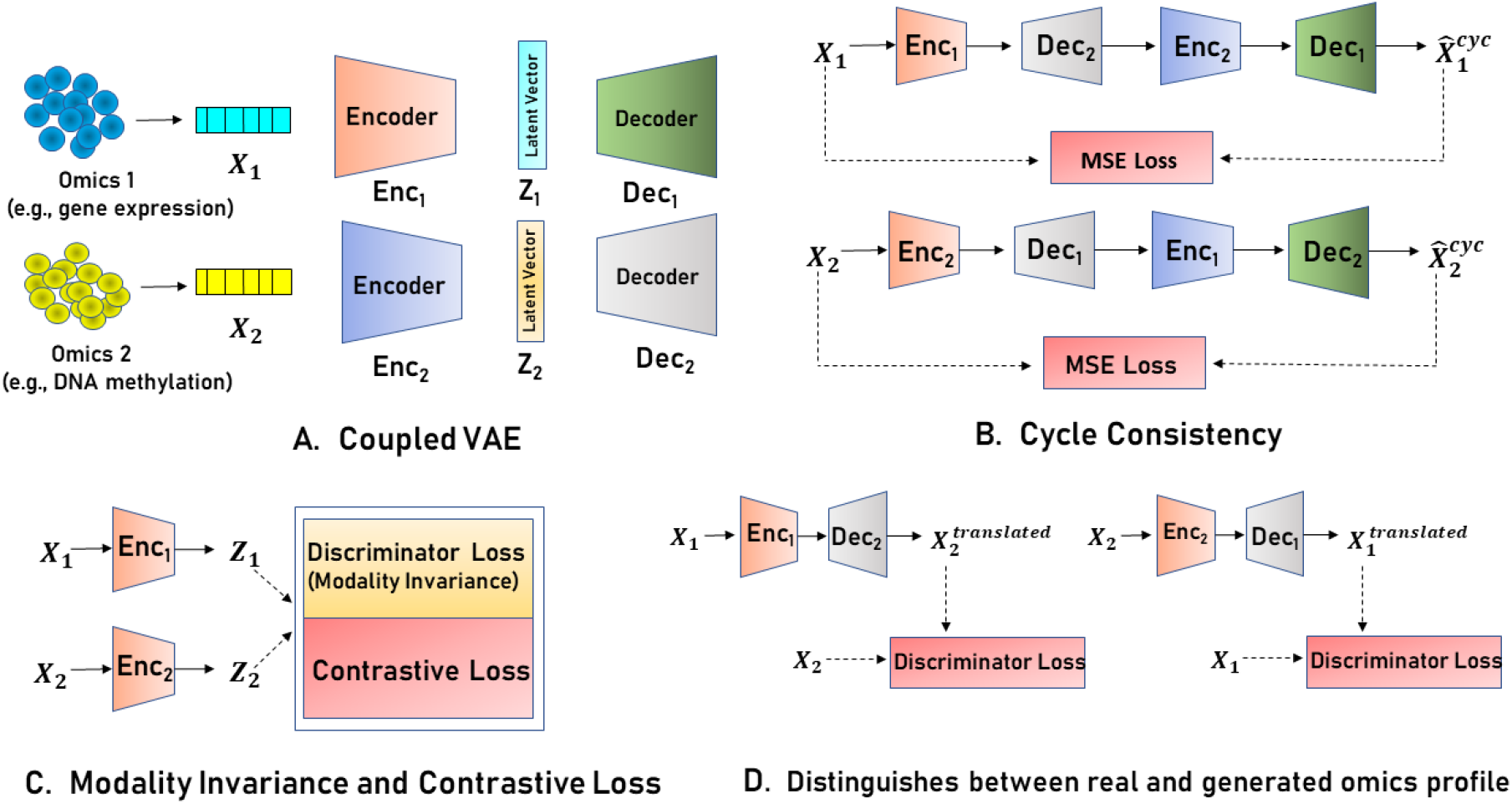
A. Coupled VAE architecture that takes two different types of unpaired omics vectors denoted as *X*_1_ and *X*_2_. B. Cycle Consistency Mechanism. C. Latent representations of both omics are trained to be modality invariant through a discriminator loss and an additional contrastive loss pushes the model to learn AD-specific patterns. D. Two additional discriminators attempt to distinguish between generated and real omics profiles, while the model aims to “fool” these discriminators, ensuring the generated profiles appear biologically realistic.

#### Variational Autoencoder (VAE)

Each VAE models the latent variable distributions for the two omics types. The encoder for each VAE approximates the posterior distribution of the latent variable **z** given the input data **x**. Specifically, for omics type **x**_1_, the encoder 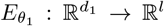 approximates 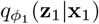, where *l* is the dimensionality of the latent space. The posterior is assumed to be a multivariate Gaussian distribution with a diagonal covariance structure: 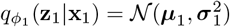 where the encoder outputs ***µ***_1_ (the mean) and 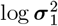 (the log-variance). Similarly, for omics type **x**_2_, the encoder 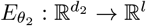 approximates 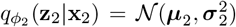 To sample from the latent distribution, we employ the reparameterization trick, which ensures differentiability by expressing **z** as a function of the mean, variance, and a noise term *ϵ* drawn from a standard normal distribution: **z**_1_ = ***µ***_1_ + *σ*_1_ · *ϵ, ϵ* ∼ 𝒩 (0, *I*), where 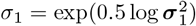 Similarly, for **z**_2_: **z**_2_ = ***µ***_2_ + *σ*_2_ · *ϵ, ϵ* ∼ 𝒩 (0, *I*), where 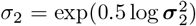. The latent variables **z**_1_ and **z**_2_ are then decoded by 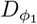 and 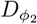 to reconstruct the input omics data:

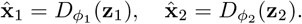

Here, 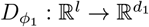 and 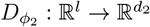 represent the decoders that reconstruct the input. To ensure that the learned posterior 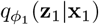 is close to the prior *p*(**z**_1_) = 𝒩 (0, *I*), a KL divergence regularizer is applied. The KL divergence for each VAE is computed as:

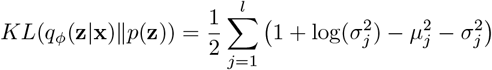

**Cycle Consistency Loss :** Without having a direct paired mapping between the two modalities, the model might fail to properly align the features of one omics type with the other. To overcome this limitation, we incorporate a **cycle consistency loss** to enforce bidirectional mapping between **x**_1_ and **x**_2_. We sample **x**_1_ ∼ *P*_data1_(**x**_1_), encode it to obtain **z**_1_, and then decode **z**_1_ into the other modality as : 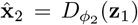 The translated data 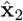 is re-encoded using 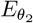 to obtain 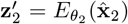 which is then decoded back to the original modality:

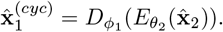

This process ensures that **x**_1_ can be recovered through the translation cycle. Similarly, the same process is applied to **x**_2_. The **cycle consistency loss** is formulated as:

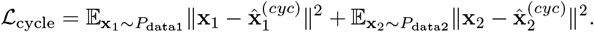

This loss encourages consistency between the two omics types, ensuring that even though the data is unpaired, the model can translate accurately in both directions and preserve the essential features of the original data. By enforcing this cycle consistency, we mitigate drift between the two modalities and effectively push the model to learn to generate paired data.

#### Adversarial Loss

In addition to cycle consistency, we utilize three adversarial discriminators to ensure modality invariance in the latent space and to enforce that the generated omics profiles are realistic. The first discriminator, *D*_*ψ*_, operates in the latent space to ensure that **z**_1_ and **z**_2_ are indistinguishable, enforcing modality invariance. Modality invariance in latent space is crucial because we use the latent space of one omics type to reconstruct the other omics profile through the opposite decoder. The adversarial loss for modality invariance is:

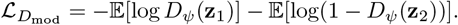

The second and third discriminators, 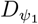 and 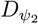, ensure that the generated omics data from 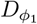 and 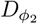, respectively, are indistinguishable from real data. For **x**_1_, the adversarial loss is given by:

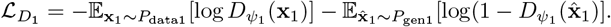

Similarly, for **x**_2_, the adversarial loss is:

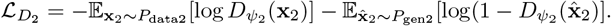

*P*_gen1_ and *P*_gen2_ represent the distributions of the generated omics profiles from the cross-omics translation. These adversarial losses ensure that the generated omics profiles remain biologically plausible and realistic, and that the latent space enforces modality invariance.

#### Contrastive Loss

To capture AD-specific features in the latent space, we employ a **contrastive loss** that encourages latent representations of omics data from the same class (AD or control) to be closer in the latent space. For a pair of latent variables **z**_1_ and **z**_2_ which represent positive pairs meaning samples with the same label *y*_1_ = *y*_2_ ∈ {0, 1}, the contrastive loss is defined as:

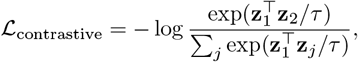

where *τ* is a temperature parameter that controls the concentration of the distribution and *j* runs over all samples in the batch, including both positive and negative samples.

#### Overall Objective

The total loss for this translation module is defined as :

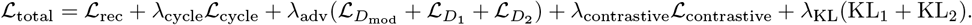

where 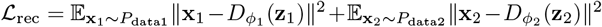 and lambdas control the relative importance of each loss component. This formulation ensures that the translated omics data remains biologically realistic, AD-specific features are captured, and the unpaired nature of the data is properly handled through cycle consistency.

### 2.2 Prediction Module

Let 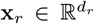 represent the real omics profile of one type, and 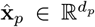 represent the predicted omics profile of another type, generated by the pretrained translation module. *d*_*r*_ and *d*_*p*_ represent the respective dimensionalities of the real and predicted omics types. The pretrained Cross-Omics Translation Module *T* maps the real omics profile to the predicted omics profile as follows: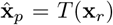, where *T* is the mapping function learned during the translation training phase. The prediction module combines these two profiles, 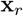 and 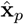, to perform AD classification (Figure 2). The fusion of the real and predicted omics profiles is performed by projecting each profile into a lower dimensional space of same dimension. The real omics profile **x**_*r*_ and the predicted omics profile 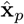 are separately projected using linear transformations: **g**_*r*_ = **W**_*r*_**x**_*r*_ and 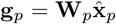, where 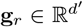 and 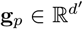 are the projected representations, and 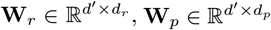 are learnable projection matrices. To dynamically weigh the contributions of each omics profile type, we apply learnable element-wise weights **m**_*r*_ and **m**_*p*_ : **h**_*r*_ = **g**_*r*_ ⊙**m**_*r*_ and **h**_*p*_ = **g**_*p*_ ⊙**m**_*p*_, where ⊙denotes element-wise multiplication. Both **m**_*r*_ and **m**_*p*_ are of same dimension *d’*^′^.The fused latent representation is then computed as:

**Fig. 2:**
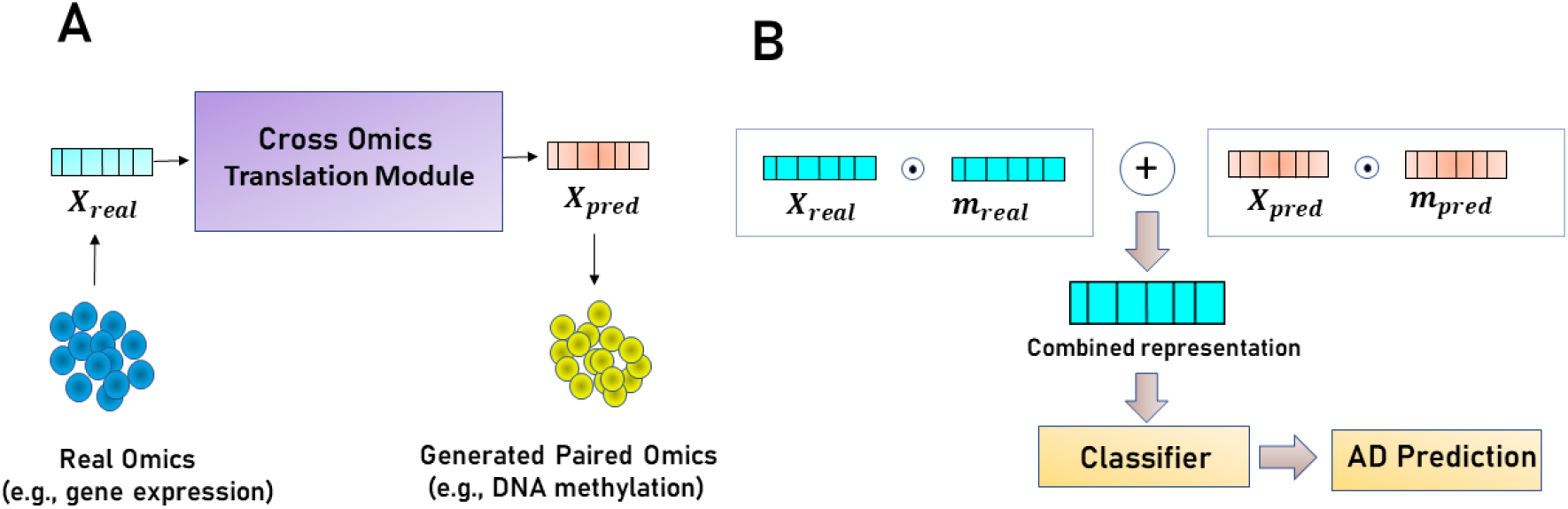
A. Cross Omics Translation Module generates corresponding paired omics profile (e.g, gene expression) from real omics profile (e.g, DNA methylation or proteomics). B. Architecture of prediction module. *X*_*real*_ and *X*_*pred*_ are feature vectors of the real and generated omics and *m*_*real*_ and *m*_*pred*_ are learnable weight vectors. ⊙ is element-wise multiplication.

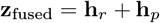

This fused representation 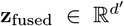 captures complementary information from both the real and predicted omics profiles of different types, and it will be used for AD classification. The fused representation **z**_fused_ is passed through a multi-layer classifier to predict the probability of Alzheimer’s Disease. The classifier is composed of multiple fully connected layers with ReLU activations and batch normalization to ensure robust learning. The classifier computes the prediction as follows: *ŷ* = *σ*(𝒞 (**z**_fused_)), where is the classifier’s transformation and *σ*(·) is the sigmoid activation function, which outputs *ŷ* ∈ [0, 1]. The prediction module is trained to minimize the binary cross-entropy loss between the predicted probability *ŷ* and the true label *y* ∈ {0, 1} for Alzheimer’s Disease.

## 3 Experiments

We validated the performance and effectiveness of UnCOT-AD by conducting different experiments. We divided the experiments in two parts - one is AD prediction performance by integrating multi-omics data and the other is cross-omics translation performance analysis. For the experiments, we consider three different types of omics - gene expression, DNA methylation and proteomics. We performed 5 fold cross validation for each of the experiments.

### 3.1 Dataset

We collected the preprocessed gene expression and DNA methylation dataset from [6] and proteomics dataset from [17]. The gene expression data were obtained by integrating GSE33000 [18] and GSE44770 [2], containing 257 normal and 439 AD samples. DNA methylation data were collected from GSE80970 [19], comprising 68 normal and 74 AD samples. Proteomics dataset is comprised of 328 AD and 91 normal samples. Differentially expressed genes (DEGs) and differentially methylated positions (DMPs) were identified by filtering with P-value *<* 0.01 for DEGs, and P-value *<* 0.01 for DMPs and P-value *<* 0.05 for differentially expressed proteins(DEPs). Finally we got 200 DEGs, 500 DMPs and 696 DEPs. Detailed preprocessing steps can be found in [6,17].

### 3.2 UnCOT-AD Improves AD Prediction by Integrating Multi-Omics Data Over Single Omics

Initially, we trained individual classifiers on three separate omics datasets (gene expression, DNA methylation, and proteomics) to predict Alzheimer’s Disease (AD). Next, we trained three Cross-Omics Translation Modules: one for gene-proteomics translation, one for gene-DNA methylation translation, and another for DNA methylation-proteomics translation, all in a bidirectional manner. After training the translation modules, we generated predictions by combining real and translated omics data. For example, we took the real gene expression data and used the pretrained gene-to-proteomics translation module to generate the corresponding proteomics profile. The real gene expression data and the predicted proteomics profile were then passed into the prediction module for AD prediction. We repeated this process for all possible pairs of three different omics.

Table 1 presents the results of AD prediction using both single omics and integrated multi-omics data. The performance of gene expression and proteomics, with accuracies of 0.8765 and 0.8904 respectively, suggests that these modalities offer substantial predictive power when used alone. However, DNA methylation, with an accuracy of 0.8335, performs less effectively on its own, indicating that it may not capture the full complexity of AD on its own. Despite these reasonable results for single omics datasets, relying on a single data source limits the ability to fully capture the diverse biological signals related to AD.

**Table 1:** Performance Metrics for AD Prediction.

| Omics Type | Accuracy | Precision | Recall | F1 Score | MCC |
| --- | --- | --- | --- | --- | --- |
| **Multi Omics** |  |  |  |  |  |
| Gene+Predicted DNA Methylation | **0.9498** | 0.9743 | 0.9453 | **0.9594** | **0.8950** |
| Gene+Predicted Protein | 0.9427 | 0.9662 | 0.9429 | 0.9540 | 0.8792 |
| Protein+Predicted Gene | 0.9189 | 0.9487 | **0.9484** | 0.9474 | 0.7759 |
| Protein+Predicted DNA Methylation | 0.9429 | **0.9958** | 0.9303 | 0.9587 | 0.8842 |
| DNA Methylation+Predicted Protein | 0.8828 | 0.8857 | 0.8800 | 0.8828 | 0.7657 |
| DNA Methylation+Predicted Gene | 0.8759 | 0.8842 | 0.9067 | 0.8941 | 0.7469 |
| **Single Omics** |  |  |  |  |  |
| Gene | 0.8765 | 0.8922 | 0.9157 | 0.9035 | 0.7335 |
| DNA Methylation | 0.8335 | 0.8312 | 0.8667 | 0.8474 | 0.6674 |
| Protein | 0.8904 | 0.9703 | 0.8879 | 0.9232 | 0.7636 |

The integration of multi-omics data provides both notable quantitative improvements and biological insights into AD prediction. Combining gene expression with predicted DNA methylation achieves an 8.4% accuracy improvement over gene expression alone and a 13.9% improvement over DNA methylation alone, suggesting that transcriptional activity coupled with epigenetic information provides better understanding of AD mechanisms. Likewise, gene expression combined with predicted proteomics yields a 7.5% improvement over gene expression alone and 5.9% over proteomics alone, highlighting the complementary roles of transcriptional and protein-level data. Protein expression with predicted gene expression reaches an accuracy of 0.9189, underscoring how upstream genetic regulation benefits protein-level prediction. Integrating protein expression with predicted DNA methylation or DNA methylation with predicted proteomics improves accuracy by 5.9% and 5.1%, respectively, reinforcing that combining epigenetic information with proteomics leads to a more comprehensive model for AD prediction. We also report the Matthews Correlation Coefficient (MCC) to provide a more comprehensive evaluation of the model’s performance [20]. For binary classification tasks, MCC is particularly useful as it takes into account the balance between true positives, false positives, true negatives, and false negatives. The highest MCC value is observed for the integration of gene expression with predicted DNA methylation (MCC of 0.8950) and the next is protein with predicted DNA methylation (MCC of 0.8842). These results indicate that the multi-omics models perform robustly across all confusion matrix categories. We observe that the trend of improvement after integrating multi-omics data over single omics is consistent accross all pairs of omics in every evaluation metric.

### 3.3 AD Prediction Performance Comparison with Baseline Models

We compared the performance of UnCOT-AD on multi-omics based AD prediction with three baseline models: Abbas et al., Mahendran et al., and Park et al.[12,13,6]. All of them used gene expression and DNA methylation data. So, we trained their models with proteomics data as well and conducted a thorough performance comparison. However, none of the baseline models utilized paired omics data. They used all possible pairs of gene expression and DNA methylation profile for each label - normal and AD. We address this limitation in UnCOT-AD by cross omics translation which gives us paired data. To show the effectiveness of UnCOT-AD, we compared the baseline’s performance using gene expression and DNA methylation with our results from both gene expression + predicted DNA methylation and DNA methylation + predicted gene expression. From Table 2, we observe that UnCOT-AD gives better performance in both cases compared to the baseline’s performance. This allowed us to show the added value of our cross-omics translation method. Similarly, we extended this comparison across all possible pairs of omics. We observe that UnCOT-AD consistently outperforms the baseline models across all evaluation metrics. The key factor behind this improvement is the use of our cross-omics translation module, which effectively generates paired omics data from unpaired datasets.

**Table 2:**
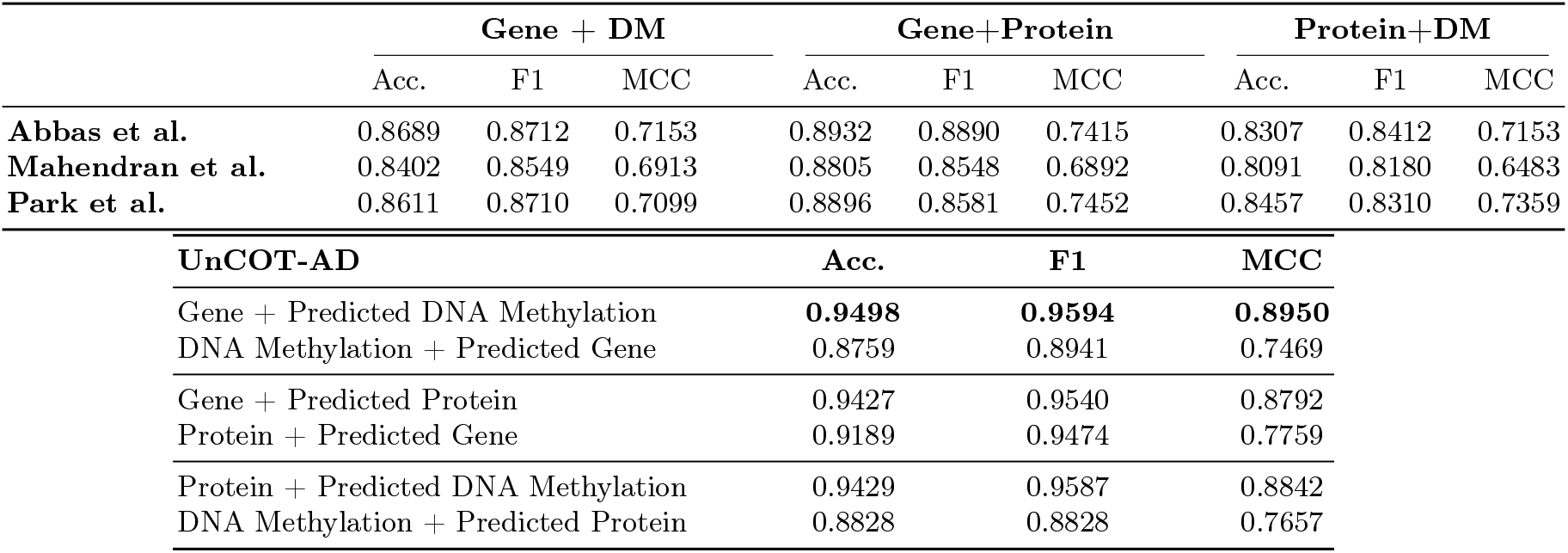
Comparison of UnCOT-AD with Baseline Models on Multi-Omics AD Prediction.

### 3.4 UnCOT-AD Captures Alzheimer’s Disease-Specific Patterns

To validate that UnCOT-AD successfully captures Alzheimer’s Disease-specific patterns, we take a real omics profile (e.g., gene expression) and use the cross-omics translation module to generate the corresponding omics profile (e.g., DNA methylation or proteomics). The generated profile is then passed through its respective encoder, and the resulting latent representations are plotted using t-SNE. From Figure 3, we observe that AD and normal samples are well-separated on the latent space, as intended by the contrastive loss used during training. This approach shows us UnCOT-AD effectively captures disease-specific patterns in the latent space. However, we notice that the ones generated from DNA methylation (Figure 3d,3f) struggle to capture AD specific features resulting in more sparsed distribution of AD and normal samples. On the other hand, the ones generated from gene expression and proteomics show well clustured AD and normal samples.

**Fig. 3:**
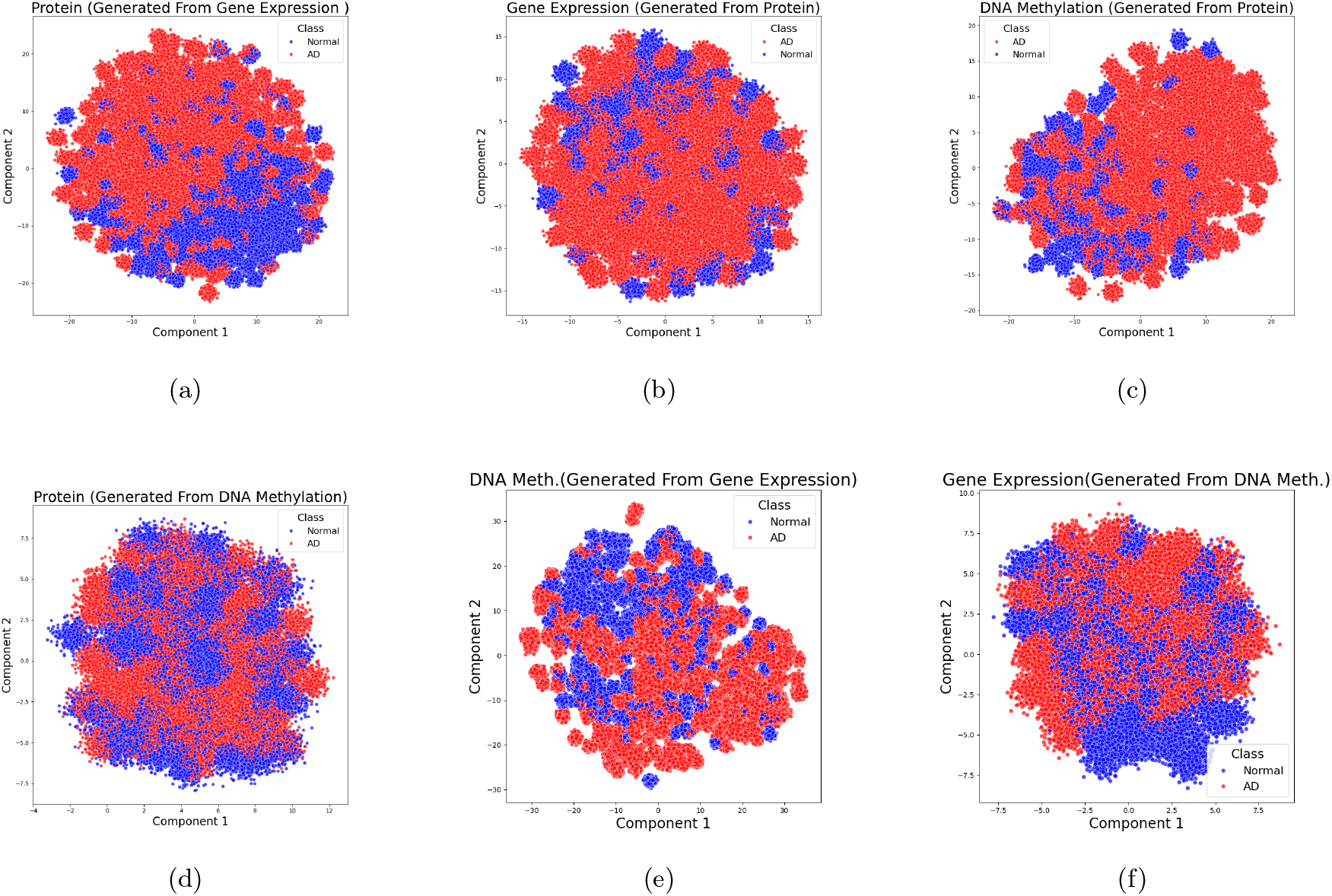
t-SNE visualization of translated omics profiles generated from real omics, with AD and control samples shown in different colors.

### 3.5 Cross Omics Translation Performance

To assess the quality of cross-omics translation, researchers commonly use correlation-based metrics such as Pearson or Spearman correlations. However, these methods are typically applied in paired omics translation, where ground truth for cross-omics data is available for each sample. In our unpaired scenerio, where no such ground truth exists for the translated omics profiles, correlation-based evaluation is not applicable. To address this challenge and effectively assess the performance of unpaired cross-omics translation, we propose two evaluation metrics: Cycle Consistency Loss (CCL) and Fréchet Omics Distance (FOD). CCL measures how well the original omics data can be reconstructed after cross-translation between different omics modalities, serving as a key indicator of the preservation of relevant features during translation. A low CCL indicates that the translated omics profile successfully preserves the underlying structure of the original data, suggesting that the translation process generates omics data closely aligned with, or effectively “paired” to, the original modality. A lower FOD quantifies the lower distributional difference between real-world omics data and the omics profile generated through cross-omics translation, capturing how closely the generated omics data aligns with the real omics profile in the latent space.

#### Cycle Consistency Loss (CCL)

The Cycle Consistency Loss (CCL) of *X*_1_ to *X*_2_ (*X*_1_ –> *X*_2_) is computed as :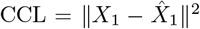 . where *X*_1_ represents the input omics data from modality 1. First, *X*_1_ is passed through the encoder Enc_1_ of modality 1 to obtain its latent representation. This is then decoded using the decoder Dec_2_ of modality 2 to generate the predicted omics 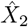. The predicted omics 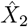 is passed through the encoder Enc_2_, followed by the decoder Dec_1_, resulting in the reconstructed data 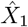. The CCL is then calculated as the mean squared error (MSE) between the original *X*_1_ and the reconstructed 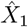.

#### Fréchet Omics Distance (FOD)

Given two omics *X*_1_ and *X*_2_, let 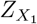 represent the latent space of *X*_1_ obtained via the encoder of the VAE corresponding to *X*_1_, denoted as 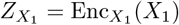. The translated omics 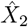 (from *X*_1_) is generated by decoding 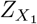 using the decoder of the VAE corresponding to *X*_2_, denoted 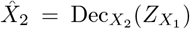. The real omics *X*_2_ and The translated omics 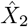 are both passed through the encoder of the VAE of *X*_2_, denoted as 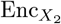, to obtain their respective latent representations 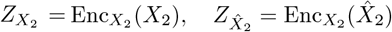. The Fréchet Omics Distance (FOD) between *X*_2_ and 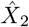 is then computed

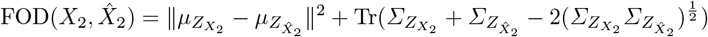

where 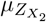 and 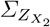 are the mean and covariance of the latent representations of real *X*_2_, and 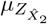 and 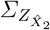 are the corresponding statistics for the predicted 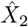.

#### Performance Analysis

We compared the bidirectional cross omics translation between gene expression, DNA methylation and proteomics. As baselines, we employed CycleGAN (adapted for omics data) [21], an autoencoder based architecture where we replace VAE with autoencoder, and a random baseline. From Table 3, we observe that UnCOT-AD significantly outperforms all the baselines for all pairs of omics which shows the effectiveness of our approach. Moreover, we notice the lowest CCL for translation between gene expression and proteomics. The cross modal translation involving DNA methylation with both gene expression and protein demonstrates higher CCL compared to gene and protein. This same pattern is observed with FOD as well where cross modal translation involving DNA methylation yields higher FOD compared to gene and protein. These results highlight the strong biological relationship between gene expression and proteins and the challenge of capturing complex pattern of DNA methylation. From Section 3.2, we also notice that DNA methylation shows less predictive power compared to the other two which aligns with the trend observed in this experiment.

**Table 3:** Performance on Unpaired Cross Omics Translation.

|  | UnCOT-AD |  | CycleGAN |  | Autoencoder |  | Random Baseline |  |
| --- | --- | --- | --- | --- | --- | --- | --- | --- |
|  | CCL | FOD | CCL | FOD | CCL | FOD | CCL | FOD |
| Gene → Protein | 0.0476 | **0.5404** | 0.1254 | 1.2543 | 0.1804 | 2.2015 | 3.2831 | 4.7534 |
| Protein → Gene | **0.0396** | 1.3961 | 0.1209 | 2.1543 | 0.1754 | 2.8054 | 2.9103 | 3.6742 |
| Gene → DM | 0.1509 | 4.2545 | 0.2304 | 5.1230 | 0.2800 | 5.8056 | 4.5702 | 6.1223 |
| DM → Gene | 0.1070 | 4.0135 | 0.1953 | 4.8502 | 0.2456 | 5.6743 | 4.1209 | 6.0054 |
| DM → Protein | 0.1169 | 4.1921 | 0.2105 | 5.1344 | 0.2603 | 5.9801 | 4.3658 | 6.2917 |
| Protein → DM | 0.1513 | 3.8759 | 0.2356 | 4.7523 | 0.2901 | 5.3624 | 4.4503 | 5.8421 |

### 3.6 UnCOT-AD Shows Superior Performance in Paired Cross-Omics Translation

To further demonstrate the effectiveness of UnCOT-AD, we evaluated its performance in traditional paired scenario with two state-of-the-art models, BABEL [14] and Polarbear [15], both designed specifically for paired data training. We evaluated on two paired gene expression and proteomics datasets, which were originally sourced from the TCGA BRCA cohort and made available in preprocessed form by [22]. The evaluation was conducted based on Pearson correlation between real and predicted omics profiles. UnCOT-AD consistently outperformed both BABEL and Polarbear across translation tasks in both directions: gene-to-protein and protein-to-gene (Figure 4). This result highlights UnCOT-AD’s robustness and generalizability in both paired and unpaired training scenarios.

**Fig. 4:**
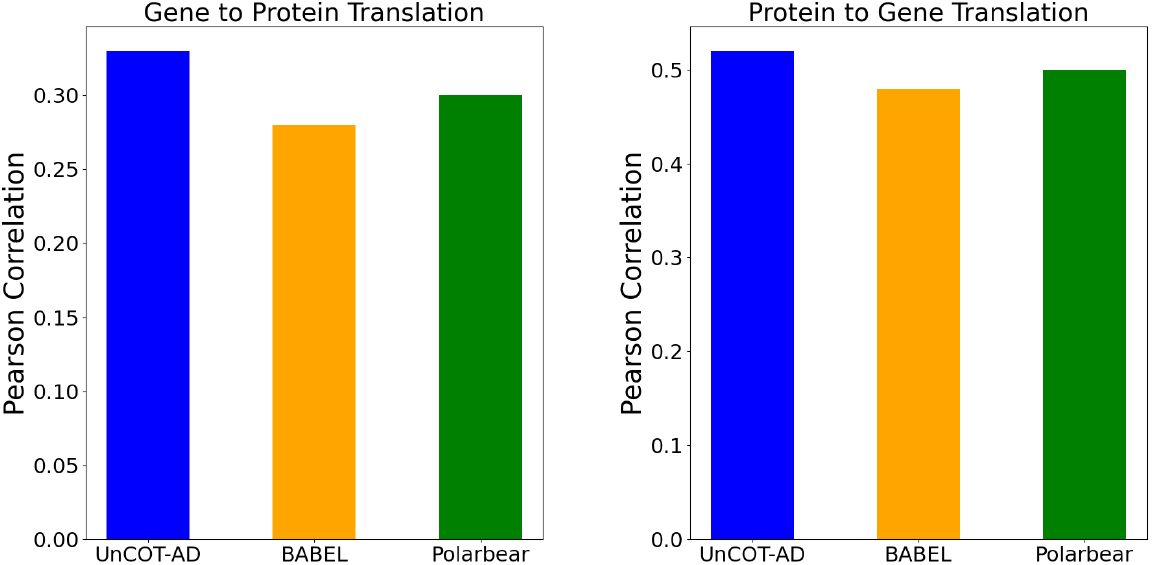
Comparison of Pearson correlation between real and predicted omics profiles across UnCOT-AD, BABEL, and Polarbear for gene-to-protein and protein-to-gene translations.

### 3.7 Ablation Study

We performed an ablation study to evaluate the impact of key components in our framework: the Adversarial Loss and Cycle Consistency Mechanism on cross-omics translation, and the contribution of the Contrastive Loss in AD prediction. From Figure 5a, we observe that removing the cycle consistency mechanism significantly increases the CCL while affecting the FOD to a lesser extent. This is because the cycle consistency mechanism is crucial for generating omics profiles that can accurately reconstruct the input omics. The absence of this mechanism leads to poor alignment between the translated and original omics profiles, increasing the CCL. On the other hand, FOD is more related to generating realistic omics profiles. Thus, removing the adversarial loss leads to substantial increase in FOD, as the adversarial components helps ensure the generated omics data resemble the true distribution of omics profiles. Therefore, to achieve optimal performance in both metrics (CCL and FOD), both the cycle consistency mechanism and adversarial loss are necessary. Additionally, we validated the importance of learning AD-specific features in the cross-omics translation module. From Figure 5b, we find that when the contrastive loss is excluded, the MCC of AD prediction drops for every pair of omics data. This highlights the essential role of contrastive loss in encouraging the separation of AD and control samples in the latent space.

**Fig. 5:**
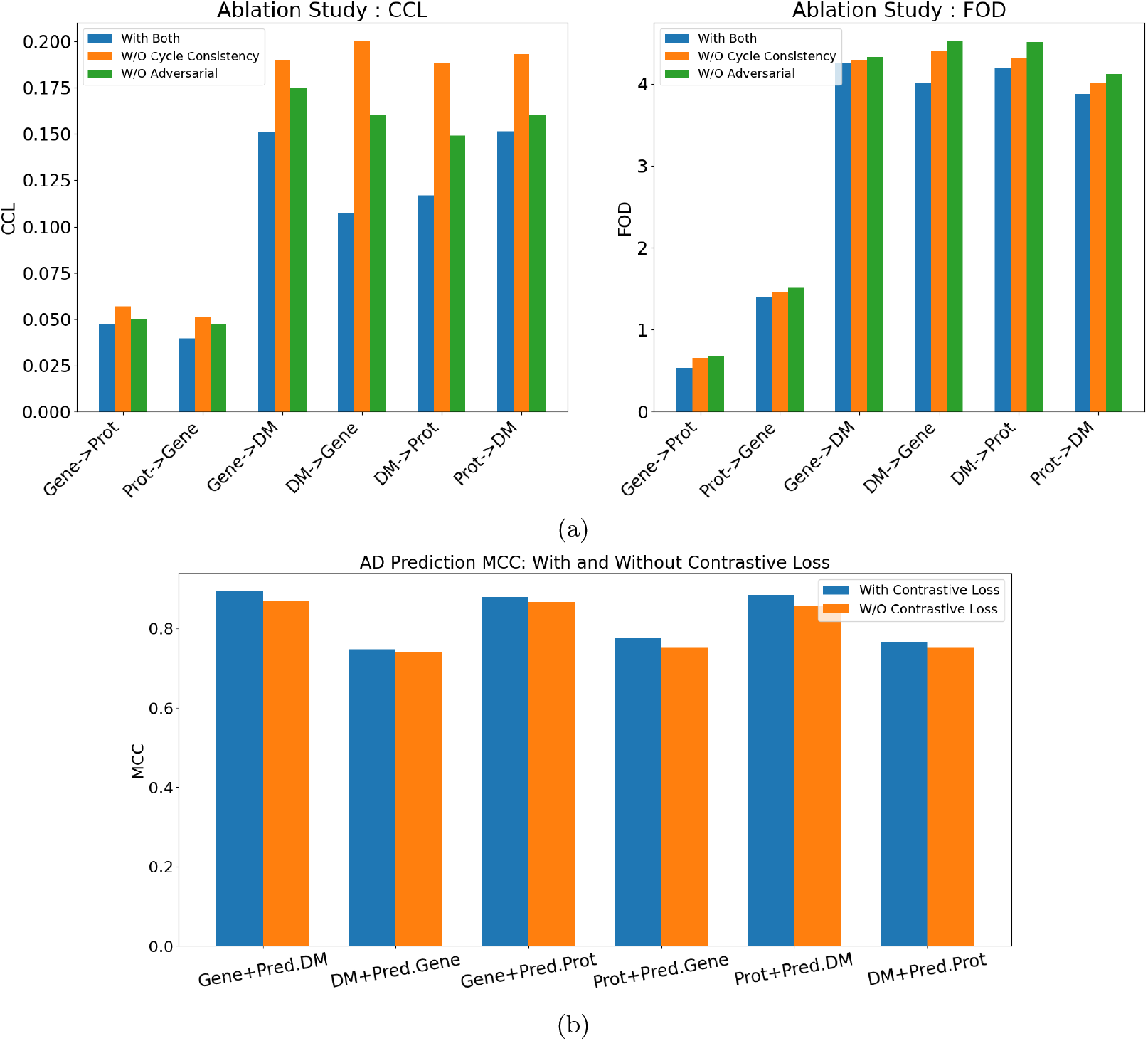
Visualization of the Contribution of (a) Adversarial Loss, Cycle Consistency and (b) Contrastive Loss in Cross Omics Translation and AD Prediction.

## 4 Conclusion

In this paper, we introduce UnCOT-AD, a novel approach to address the challenge of unpaired multi-omics data integration for Alzheimer’s Disease (AD) prediction. For the first time, we introduce a cross-omics translation module that allows for the generation of paired omics profiles, even when the training data from two modalities are unpaired. By combining the generated profiles with real omics data, we perform multi-omics based AD prediction, overcoming the limitations of existing approaches that rely solely on single omics. Our method achieves state-of-the-art results in both cross-omics translation and AD prediction tasks, demonstrating significant improvements in all evaluation metrics compared to existing approaches. By generating biologically meaningful omics profiles and effectively integrating them, our framework has proven to be a robust solution for multi-omics integration in AD prediction. Thus, UnCOT-AD presents new potential by effectively combining multiple biological data modalities, enabling a more comprehensive understanding of complex diseases like Alzheimer’s Disease. Our cross-omics translation module is designed to be compatible with any two omics types. Additionally, as our translation and prediction modules are separate, our method can be applied solely for cross-omics translation between two modalities, even in the absence of paired data.

## Notes

### Competing Interest Statement

The authors have declared no competing interest.

